## Supplementary figures and legends for "Transcription factor paralogs orchestrate alternative gene regulatory networks by context-dependent cooperation with multiple cofactors"

a

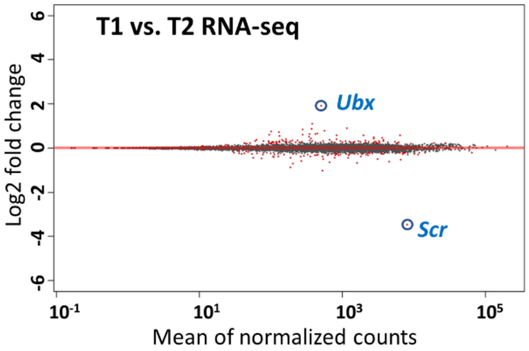


b

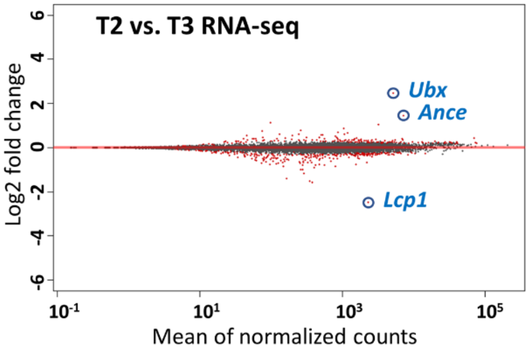


c

- FDR<0.01 AND abs(log2(fold change))>1

| | up | | | |
|---|---|---|---|---|
| down | | T1 | T2 | T3 |
| | T1 | | 2 | 8 |
| | T2 | 2 | | 4 |
| | T3 | 20 | 6 | |

Extended Data Fig. 1

**Extended Data Fig. 1. The transcriptomes of the 3 pairs of leg discs are highly similar**

**a.** MA plot comparing the transcriptomes of T1 and T2 leg discs. Differentially expressed genes, which are defined as FDR<0.01, are labeled red.

**b.** Similar to **a**, except that the comparison is between T2 and T3 leg discs.

**c**. Table showing the number of differentially expressed genes with at least a 2-fold change for all pair-wise comparisons.

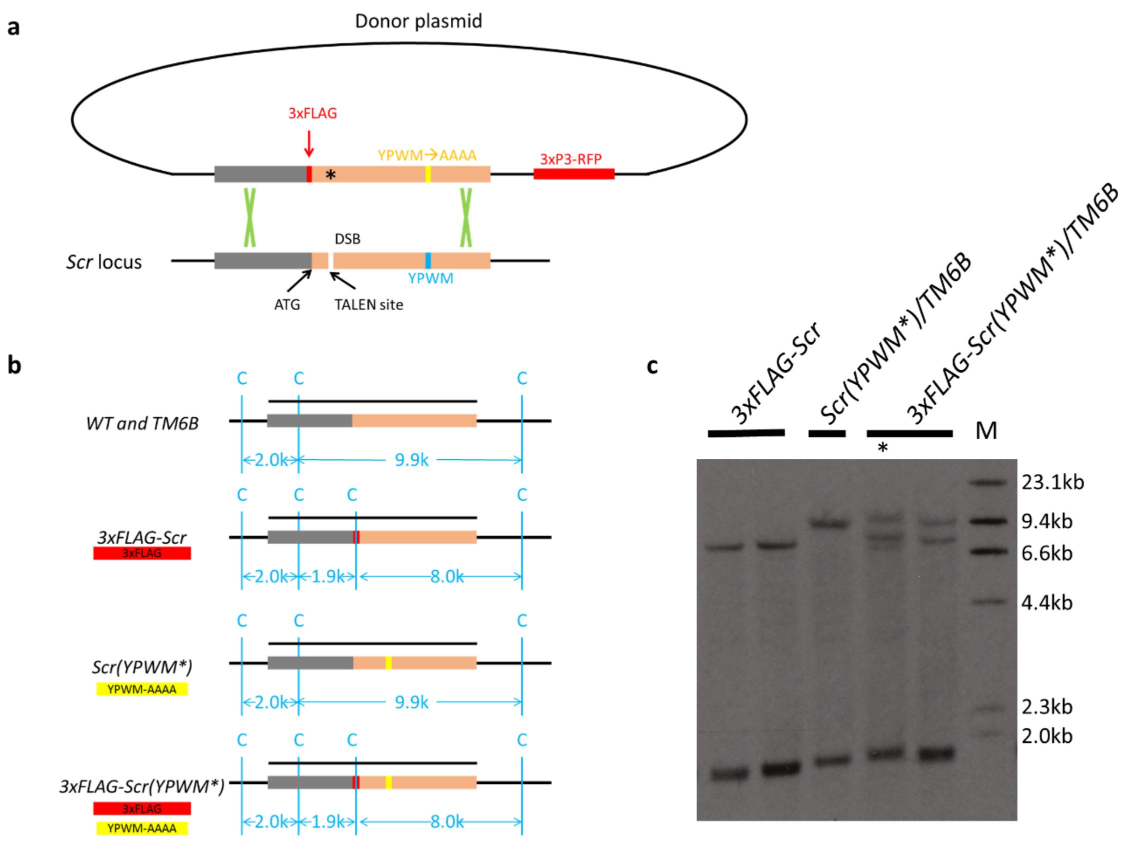


Extended Data Fig. 2

**Extended Data Fig. 2. The generation of *3xFLAG-Scr* and *3xFLAG-Scr(YPWM*)* alleles.**

**a.** The strategy used in targeting the *Scr* locus using a pair of TALENs. The orange boxes indicate coding region, and the grey boxes are the 5'UTR. The positions of the ATG start codon, 3xFLAG tag, TALEN target site, the YPWM motif, and the mutated YPWM motif on the donor plasmid are shown. The asterisk indicates that the TALEN target on the donor plasmid is mutated, so the TALENs only cut the genomic DNA. 3xP3-RFP was included on the donor plasmid to select against whole plasmid integration events (see Methods). DSB: double stranded DNA break.

**b.** ClaI restriction map of various genotypes. The blue lines indicate the ClaI sites relevant to the Southern blot. The positions of the 3xFLAG tag and the YPWM-AAAA mutation are indicated when applicable. The black bars above the genomic loci indicate the fragment used as Southern probe, and the sizes of expected Southern bands are shown.

**c.** Southern blot result of several alleles obtained in genomic targeting of the *Scr* locus. M: Roche DIG labeled DNA marker II. Asterisk: a potentially erroneous allele with a Southern blot pattern deviated from the expected one.

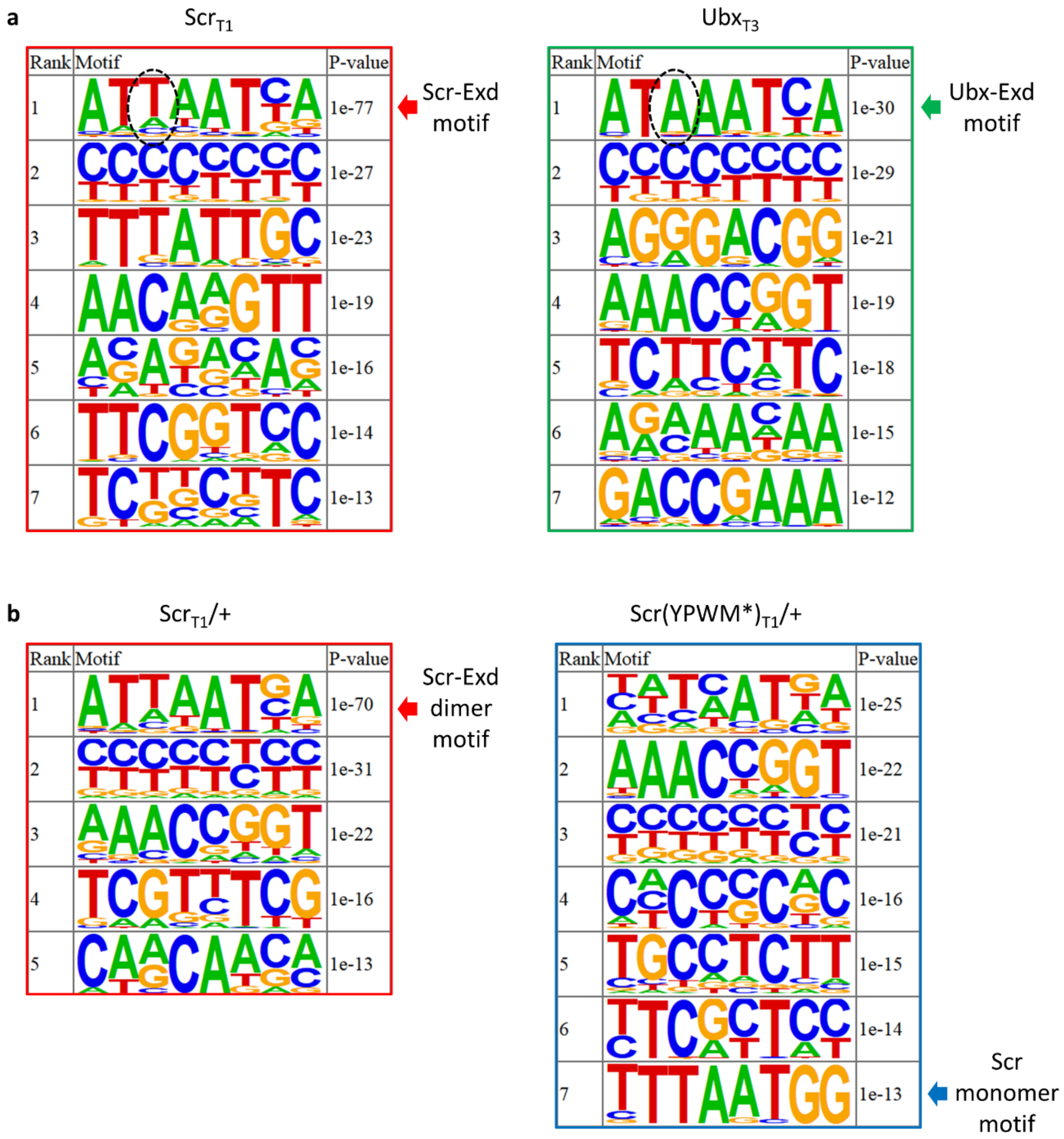


Extended Data Fig. 3

**Extended Data Fig. 3. Sequences identified by *de novo* motif searches.**

**a.** Motifs enriched in $Scr_{T1}$ and $Ubx_{T3}$ peaks that are located in intergenic or intronic regions of the genome. Arrows point out known Scr-Exd and Ubx-Exd dimer motifs. Dashed ovals highlight the positions within the Hox-Exd heterodimer motifs that Scr and Ubx are known to have different preferences [3]. Up to 7 significantly enriched motifs are shown.

**b.** Motifs enriched in $Scr_{T1}$/+ and Scr(YPWM*)$_{T1}$/+ peaks located in intergenic and intronic regions. Red arrow indicates the known Scr-Exd dimer motif, and blue arrow indicates a potential Scr monomer motif. Up to 7 significantly enriched motifs are shown.

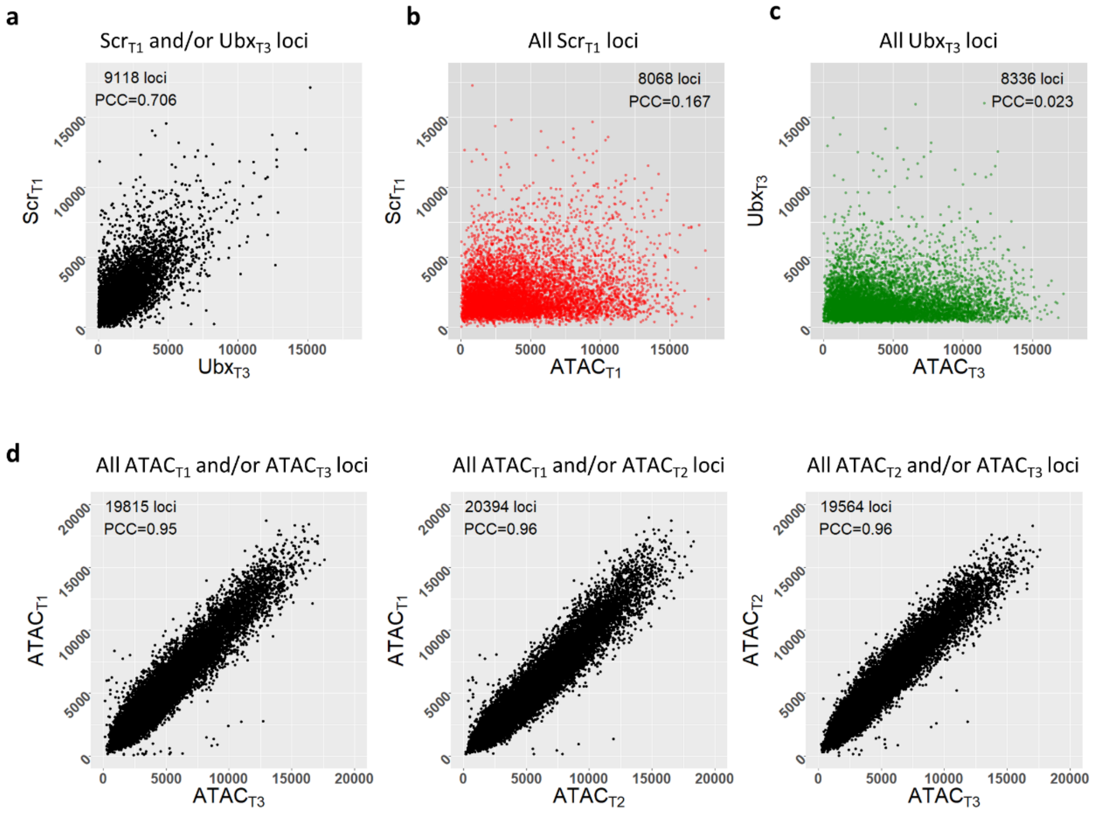


Extended Data Fig. 4

**Extended Data Fig. 4. Most paralog-specific Hox bound loci do not show tissue-specific differences in chromatin accessibility.**

**a.** Scatter plot comparing $Scr_{T1}$ and $Ubx_{T3}$ ChIP-seq signals in all Hox bound loci. Hox bound loci are defined as loci bound by either Scr in T1 leg discs or by Ubx in T3 leg discs, or by both (see Methods). The Pearson's correlation coefficient (PCC) between $Scr_{T1}$ and $Ubx_{T3}$ signals in all Hox bound loci is reported.

**b and c.** Correlation between $Scr_{T1}$ ChIP-seq and $ATAC_{T1}$ signals in all loci bound by Scr in the T1 leg discs (**b**), as well as that between $Ubx_{T3}$ ChIP-seq and $ATAC_{T3}$ signals in all loci bound by Ubx in the T3 leg discs (**c**). The Pearson's correlation coefficient (PCC) is reported.

**d.** Pair-wise comparisons of ATAC-seq signals between different leg discs. The number of loci and the Pearson's correlation coefficient (PCC) are reported.

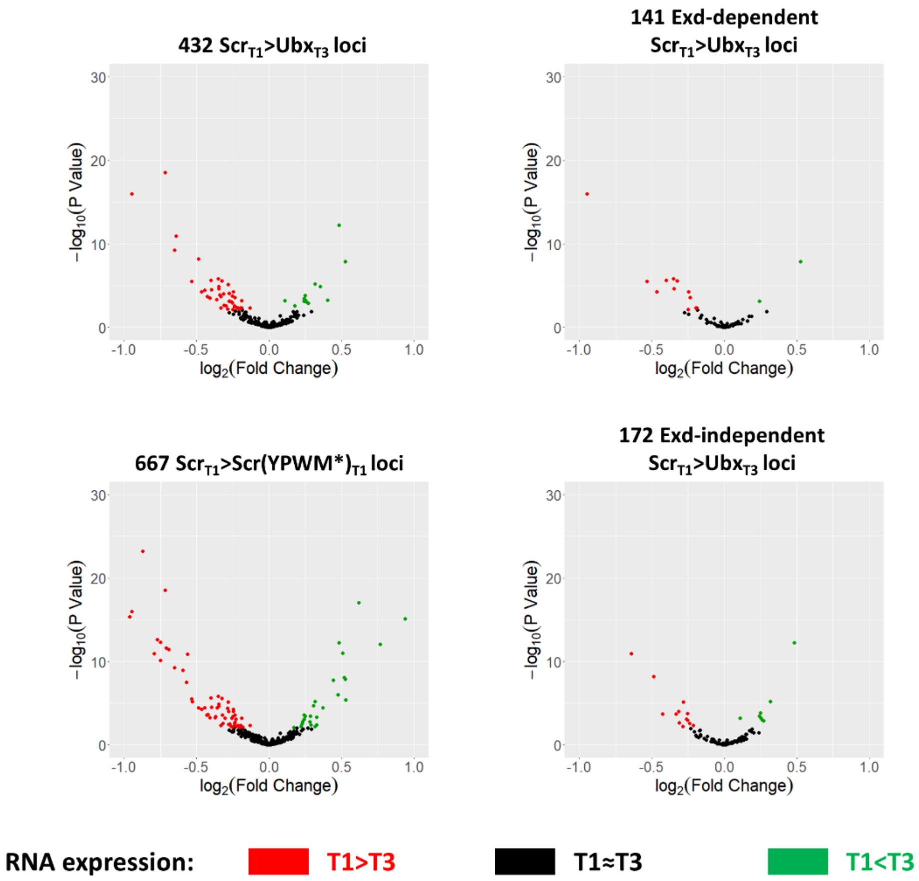


Extended Data Fig. 5

**Extended Data Fig. 5. Correlation between Hox-DNA binding and nearby gene transcription.**

Volcano plots showing T1 and T3 leg disc RNA expression of genes near different classes of Hox ChIP peaks. Genes with T1>T3 expression have negative log fold change, and those with an FDR<0.05 are colored red. Genes with T1<T3 expression have positive log fold change, and those with an FDR<0.05 are colored green.

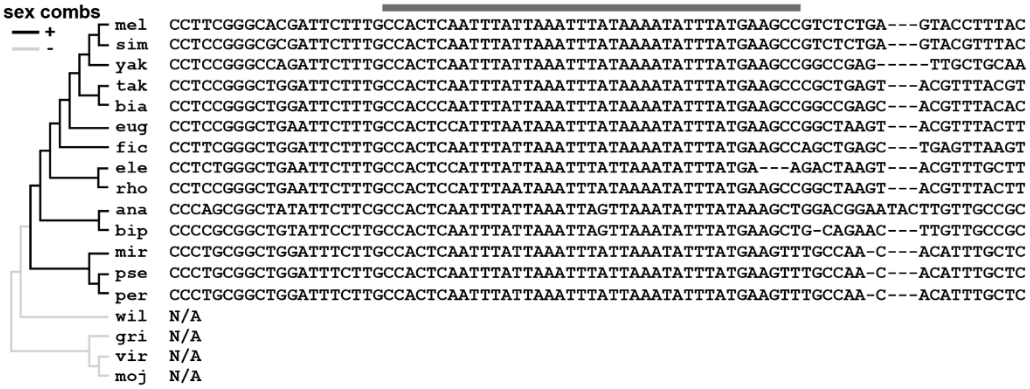


Extended Data Fig. 6

**Extended Data Fig. 6. The conservation of *dsx-1* CRM sequence correlates with the presence of sex combs.**

DNA sequence near the center of Hox ChIP peak 2 of the *dsx-1* CRM is aligned across selected *Drosophila* species. The grey bar highlights the region matches the region denoted by the vertical grey bar in Fig. 8B, and is the region of the CRM that when deleted, alters the *dsx-1* reporter gene expression. This region is highly AT-rich and contains many potential homeodomain binding sites, several of which fit the Scr-Dll consensus binding site. The presence or absence of sex comb teeth on male T1 legs is shown, and is based on [5,6].

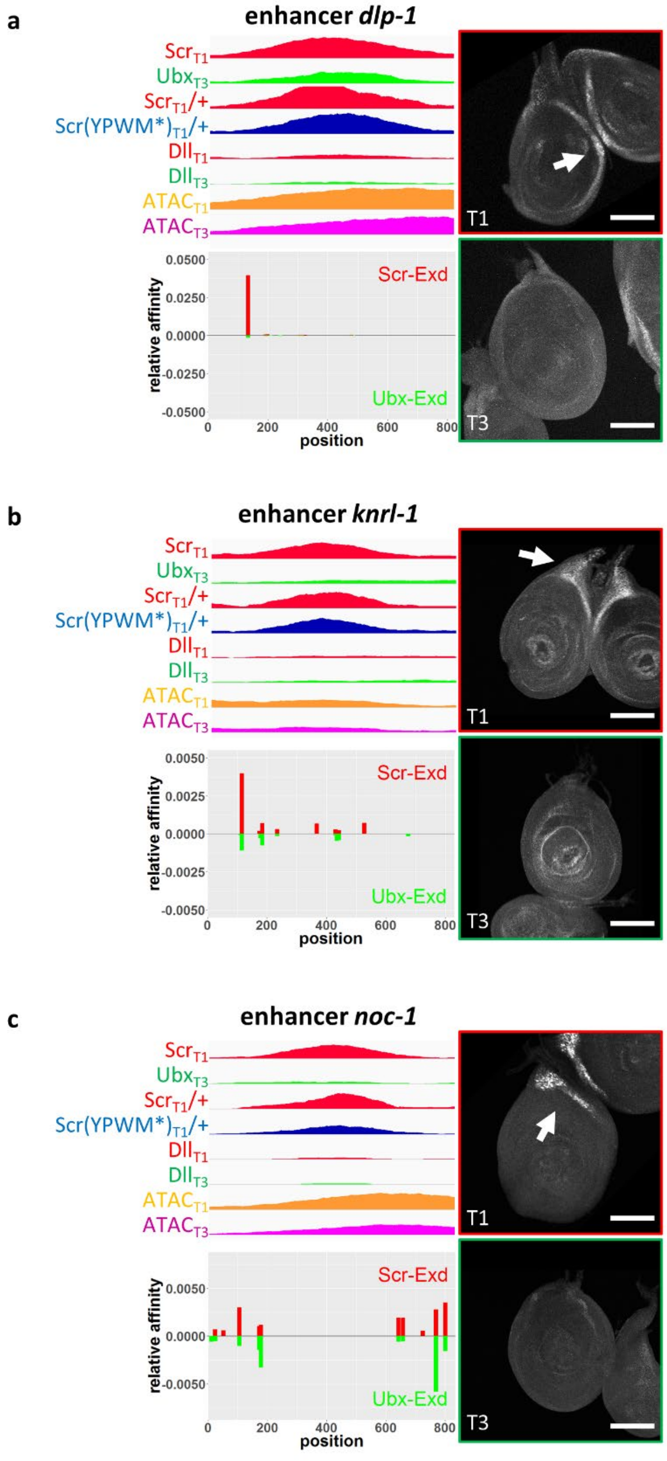


Extended Data Fig. 7

**Extended Data Fig. 7. Examples of Exd-independent $Scr_{T1}$>$Ubx_{T3}$ CRMs without Dll co-occupancy that drive T1>T3 expression**

**a.** CRM *dlp-1*. **b.** CRM *knrl-1*. **c.** CRM *noc-1*. Antibody staining results of T1 and T3 leg discs are shown, and the T1 specific expression patterns are indicated by arrows. The genome browser tracks for the $Scr_{T1}$, $Ubx_{T3}$, $Scr_{T1}$/+, Scr(YPWM*)$_{T1}$/+, $Dll_{T1}$ and $Dll_{T3}$ ChIP-seq signals, as well as $ATAC_{T1}$ and $ATAC_{T3}$ signals are shown for the genomic fragment used to generate the reporters (top left). The *NRLB* relative affinity tracks for Scr-Exd and Ubx-Exd heterodimers, aligned with the genome browser tracks, are shown at the bottom left. Note the absence of significant Hox-Exd binding motifs near Scr ChIP-seq peak center. Scale bar: 100 μm.
